# Interactome screening implicates BAG6 as a suppressor of UBQLN2 misfolding in ALS-dementia

**DOI:** 10.1101/2025.10.15.682441

**Authors:** Sang Hwa Kim, Claire E. Boos, Mark Scalf, Akasha K. Wilkemeyer, Lloyd M. Smith, Randal S. Tibbetts

## Abstract

Ubiquilin-2 (UBQLN2) is a ubiquitin (Ub)-binding shuttle protein that is mutated in X-linked forms of amyotrophic lateral sclerosis (ALS) and frontotemporal dementia (FTD). ALS/FTD-linked mutations in UBQLN2 disrupt its conformation, increasing its tendency to form cytoplasmic aggregates that may disrupt cellular regulation through loss-of-function (LOF) and gain-of-function (GOF) effects. To explore how ALS-associated mutations impact UBQLN2 function, we performed quantitative mass spectrometry (MS)-based interactome analysis using affinity-purified UBQLN2 from inducible pluripotent stem cells (iPSCs) and induced motor neurons (iMNs) expressing wild-type UBQLN2 (UBQLN2^WT^), a UBQLN2^P497H^ clinical mutant, or a UBQLN2^4XALS^ allele harboring four disease mutations. Proteins showing enhanced association with ALS-mutant UBQLN2 proteins included PEG10, a known degradation target of UBQLN2, and BAG6, a chaperone involved in the triage of mislocalized proteins (MLPs). BAG6 knockdown inhibited the solubility recovery of both wild-type and ALS-mutant UBQLN2 proteins following heat stress (HS), suggesting it functions as a UBQLN2 holdase. In addition, knockdown of BAG6 or knockout of UBQLN2 led to PEG10 accumulation, implicating both in PEG10 turnover; however, neither BAG6 nor UBQLN2 was required for PEG10 degradation in response to HS. The aggregation prone UBQLN2^4XALS^ mutant showed increased PEG10 binding and modestly delayed PEG10 turnover while PEG10 degradation was not significantly different between UBQLN2^WT^ and UBQLN2^P497H^ iPSCs. The combined findings implicate BAG6 a UBQLN2 holdase and identify a suite of proteins whose altered binding may contribute to pathologic changes in UBQLN2-associated ALS/FTD.

## Introduction

The intrinsic toxicity of misfolded, aggregation-prone proteins is normally counteracted by evolutionarily conserved proteostasis networks, which oversee the entire lifecycle of a protein— from synthesis and folding to trafficking and degradation. In aging-related neurodegenerative diseases (NDs) such as Alzheimer’s Disease (AD), Parkinson’s disease (PD), amyotrophic lateral sclerosis (ALS), and frontotemporal dementia (FTD), this delicate balance is progressively disrupted, leading to pathologic protein accumulation and neuron demise [1].

ALS is a fatal neurodegenerative condition marked by progressive degeneration of upper and lower motor neurons (MNs), resulting in muscle weakness, paralysis, and death within 3–5 years of symptom onset in the absence of mechanical ventilation [2; 3; 4]. While most ALS cases are sporadic (sALS), approximately 10–15% are familial (fALS), involving mutations in more than 40 genes. The signature histopathologic feature of ALS is the accumulation of misfolded, cytosolic TDP-43 (43 kDa TAR DNA-binding protein) which is seen in virtually all instances of ALS except those caused by mutations in superoxide dismutase 1 (SOD1) or fused in sarcoma (FUS) [5; 6]. Cytosolic aggregation and associated nuclear depletion of TDP-43 leads to cryptic exon inclusion and other RNA splicing changes in functionally important genes, triggering axonal and synaptic failure of motor neurons [7; 8; 9; 10; 11; 12; 13]. The upstream events leading to TDP-43 aggregation are poorly understood but may involve genetic susceptibility, environmental exposure, and aging-dependent reductions in proteostasis and nucleocytoplasmic transport [3; 14; 15; 16].

X-linked mutations in UBQLN2 are a rare cause of ALS, FTD, and hereditary spastic paraplegia [17; 18; 19]. UBQLN2 is a Ub-binding shuttle protein that plays a critical role in cellular proteostasis by facilitating the degradation of misfolded or ubiquitinated proteins via the proteasome and autophagy pathways. Structurally, UBQLN2 contains an N-terminal ubiquitin-like (UBL) domain and a C-terminal ubiquitin-associated (UBA) domain, connected by a methionine-rich central region that includes stress-inducible protein 1 (STI1)-like motifs [20; 21; 22; 23]. Unlike other members of the ubiquilin family, UBQLN2 is found only in eutherian mammals, suggesting a lineage-specific role, possibly including functions in placental biology [24].

ALS-associated UBQLN2 mutations cluster within a proline-rich repeat (PRR) region and are known to decrease UBQLN2 solubility, promoting cytoplasmic aggregation and interfering with its normal function in protein quality control [25; 26]. These mutations may also perturb UBQLN2 phase separation dynamics, further contributing to disease pathology. Notably, UBQLN2-positive aggregates are not restricted to genetically linked ALS cases—they are also frequently found in sALS and in ALS cases associated with *C9ORF72* repeat expansions, which are typically characterized by TDP-43 inclusions [27]. These observations suggest that UBQLN2 dysfunction may contribute to ALS pathogenesis more broadly, even in the absence of direct mutations.

Rodent models overexpressing UBQLN2^ALS^ mutants recapitulate the formation of UBQLN2 inclusions seen in ALS/FTD patients, although behavioral and neurodegenerative phenotypes vary. Transgenic mice expressing the UBQLN2^P497H^ mutation exhibit mild cognitive deficits without overt neurodegeneration, while other models show more pronounced motor neuron disease and reduced survival [28]. Interestingly, even overexpression of wild-type UBQLN2 can result in toxicity in flies and rodents, suggesting that precise regulation of UBQLN2 levels and interactions is crucial for neuronal health [29; 30; 31].

The mechanisms underlying UBQLN2-linked neurodegeneration remain incompletely understood but likely involve both LOF and GOF mechanisms driven by altered interactions and protein aggregation. To investigate this further, we conducted a quantitative proteomic screen comparing the interactomes of wild-type and ALS-associated mutant UBQLN2 in iPSC iMNs. We identified enhanced interactions of mutant UBQLN2 with PEG10—a retroelement-derived protein and established UBQLN2 degradation target [24]—and with BAG6, a molecular chaperone involved in the triage of MLPs.

Our functional studies revealed that UBQLN2 and BAG6 cooperatively regulate PEG10 levels, and that BAG6 preferentially associates with misfolded UBQLN2, acting as a holdase chaperone. Knockdown of BAG6 exacerbated UBQLN2 aggregation under proteotoxic conditions, while BAG6 overexpression suppressed UBQLN2 accumulation and toxicity. These findings establish BAG6 as a key modifier of UBQLN2 proteostasis and highlight the role of chaperone networks in buffering against UBQLN2-driven neurodegeneration.

## Materials and methods

### iPSC culture and iMN differentiation

A control human iPSC line (WC031i-5907-6; derived from neonatal male fibroblasts) was obtained from the WiCell Research Institute [32]. ALS-associated mutations were introduced into this line using CRISPR/Cas9, generating the following genotypes: P497H, 4XALS (P497H, P506T, P509S, P525S), and I498X (a 1-nt deletion at codon 497 causing a frameshift and premature stop at codon 498). All iPSC lines exhibited normal karyotypes, expressed pluripotency markers, demonstrated trilineage differentiation capacity, and were confirmed to be mycoplasma-free [32; 33]. Short tandem repeat (STR) analysis confirmed the identity, clonality, and purity of each line (95–98% confidence across 15 loci).

iPSCs were cultured on Matrigel-coated plates (Corning, 47743-720) in mTeSR1 Plus medium (STEMCELL Technologies,100-0276) and passaged every 3-4 days at a 1:3 to 1:6 split ratio using 0.5 mM EDTA and maintained at 37°C in a humidified incubator with 5% CO_2_. For the heat stress and recovery test, cells were incubated at 42°C in 5% CO_2_ for 2 hours (heat stress), followed by incubation at 37°C in 5% CO_2_ for 2 hours (recovery).

Differentiation into iMNs was performed as previously described [33; 34]). Briefly, iPSCs were dissociated and plated on Matrigel-coated plates. The following day, medium was replaced with a 1:1 mixture of DMEM/F12 and Neurobasal medium supplemented with 0.5× N2, 0.5× B27, and 1× GlutaMAX (all from Invitrogen). Small molecules were added to the medium: CHIR99021 (3 μM; Tocris), DMH-1 (2 μM; Tocris), and SB431542 (2 μM; Stemgent). Medium was refreshed every other day. After 7 days, cells differentiated into neuroepithelial progenitors (NEPs). NEPs were dissociated using 1 mg/mL Dispase and replated at a 1:6 ratio in the same neural differentiation medium supplemented with retinoic acid (RA; 0.1 μM; Stemgent), purmorphamine (0.5 μM; Stemgent), CHIR99021 (1 μM), DMH-1 (2 μM), and SB431542 (2 μM). After 7 days, cells differentiated into OLIG2^+^ motor neuron progenitors (MNPs). To initiate motor neuron differentiation, OLIG2^+^ MNPs were dissociated using 0.5 mM EDTA and cultured in suspension in neural differentiation medium supplemented with RA (0.1 μM) and purmorphamine (0.1 μM). After 6 days, cells differentiated into HB9^+^ neural motor precursors (NMPs). HB9^+^ NMPs were dissociated using Accutase (Invitrogen) and plated on Matrigel-coated plates. Cells were cultured in neural differentiation medium containing RA (0.1 μM), purmorphamine (0.1 μM), and Compound E (0.1 μM; Millipore) for 6 days to promote maturation into CHAT^+^ iMNs.

### UBQLN2 purification and mass spectrometry (MS)

To collect UBQLN2 interacting proteins, iPSCs or iMNs resuspended in a lysis buffer containing 20 mM Tris-HCl (pH 8.0), 138 mM NaCl, 10 mM KCl, 1 mM MgCl_2_, 1 mM EDTA, and 1% CHAPS (vol/vol), supplemented with 20 mM NaF, 20 mM β-glycerophosphate, and protease inhibitor cocktail. The cells were incubated on ice for 10 minutes, then centrifuged at 15,000 × g for 10 minutes at 4°C to separate soluble and insoluble fractions. The soluble fractions were immunoprecipitated with UBQLN2 antibodies with Dynabeads Protein G beads. The co-immunoprecipitated proteins were eluted with elution buffer (0.2 M Glycine, pH 2.5) and neutralized with 1 M Tris-HCl, pH 9.0. Prior to tryptic digestion, dithiothreitol (DTT) was added to a final concentration of 10 mM to reduce disulfide bonds.

UBQLN2 immunoprecipitates (IPs) were subjected to tryptic digestion using the filter aided sample preparation (FASP) method [35] and HPLC-ESI-MS/MS analysis using an Orbitrap mass spectrometer. The tryptic digest solution was desalted/concentrated using an Omix 100 µL (80 µg capacity) C18 tip. The solution was pipetted over the C18 bed 5 times, and rinsed 3 times with H_2_O, 0.1% trifluoroacetic acid (TFA) to desalt. The peptides were eluted from the C18 resin into 150 µL 70% acetonitrile, 0.1% TFA and lyophilized. The peptides were re-suspended in 95:5 H_2_O:acetonitrile, 0.2% formic acid and analyzed in duplicate as described below.

Peptides from the iPSC IPs were separated and analyzed using a UPLC-ESI-MS/MS system consisting of a NanoAcquity ultra-high-pressure liquid chromatography system (Waters) and an Orbitrap Q Exactive HF mass spectrometer (Thermo Fisher Scientific). UPLC separation employed an in-house constructed 100 x 365 μm fused silica capillary micro-column packed with 20 cm of 1.7 μm-diameter, 130 Angstrom pore size, C18 beads (Waters BEH), with an emitter tip pulled to approximately 1 μm using a laser puller (Sutter instruments). Peptides were loaded in buffer A (H_2_O, 0.2% formic acid) at a flow-rate of 400 nL/minute for 30 minutes and eluted over 120 min at a flow-rate of 300 nl/minute with a gradient of 5% to 35% acetonitrile, in 0.1% formic acid. The nano-column was held at 60°C using a column heater (in-house constructed). The nanospray source voltage was set to 2200 V. Full-mass profile scans were performed in the orbitrap between 375-1500 *m/z* at a resolution of 120,000, followed by MS/MS HCD scans of the ten highest intensity parent ions at 30% relative collision energy and 15,000 resolution, with a 2.5 m/z isolation window and a mass range starting at 100 *m/z*. Charge states 2-6 were included and dynamic exclusion was enabled with a repeat count of one over a duration of 15 seconds.

Peptides from the iMN IPs were separated and analyzed using a UPLC-ESI-MS/MS system consisting of an Easy-nLC 1200 ultra-high-pressure liquid chromatography system and an Orbitrap Fusion Lumos mass spectrometer (Thermo Fisher Scientific). UPLC separation employed an in-house constructed 100 x 365 μm fused silica capillary micro-column packed with 20 cm of 1.7 μm-diameter, 130 Angstrom pore size, C18 beads (Waters BEH), with an emitter tip pulled to approximately 1 μm using a laser puller (Sutter instruments). Peptides were loaded in buffer A (H_2_O, 0.2% formic acid) at a pressure of 300 Bar and eluted over 120 minutes at a flow rate of 350 nL/min with the following gradient established by buffer A (H_2_O, 0.2% formic acid) and buffer B (80% acetonitrile (ACN), 0.2% formic acid): Time/T = 1 min, 5% buffer B; T = 52 min, 30% buffer B; T = 80 min, 42% buffer B; T = 90 min, 55% ACN; T = 95 to 100 min, 85% buffer B; T = 101 to 120 min, equilibrate at 0% buffer B. The nano-column was held at 60°C using a column heater (in-house constructed). The nanospray source voltage was set to 2200 V. Full-mass profile scans were performed in the orbitrap between 375-1500 m/z at a resolution of 120,000, followed by MS/MS HCD scans in the orbitrap of the highest intensity parent ions in a 3 s cycle time at 30% relative collision energy and 15,000 resolution, with a 2.5 m/z isolation window. Charge states 2-6 were included and dynamic exclusion was enabled with a repeat count of one over a duration of 30 s and a 10 ppm exclusion width both low and high. The AGC target was set to “standard”, maximum inject time was set to “auto”, and 1 µscan was collected for the MS/MS orbitrap HCD scans.

### Bioinformatic analyses of mass spectrometric data

The MetaMorpheus software program (version 1.05) was used to identify peptides and proteins in the samples [36; 37]. Protein fold changes were quantified with FlashLFQ with match between runs enabled [38; 39]. The UniProt, reviewed human database (downloaded on 04/03/2024 20:34:31) as well as the MetaMorpheus contaminant database was used for calibration, global post translational modification discovery (G-PTM-D), and search. Default parameters were used including a maximum of 2 missed cleavages and minimum peptide length of 7 amino acids for a tryptic digest. Carbamidomethylation of cysteine was set as a fixed modification while oxidation of methionine was variable with 2 maximum modifications per peptide. Differential expression analysis of the quantified protein groups was performed in Perseus (version 1.6.15.0) [40]. Contaminant and decoy proteins were filtered out as well as protein groups above a protein Q-value threshold of 0.01 for the iPSC samples and 0.05 for the iMN samples. The quantified intensities were log2 transformed and normalized by Z-score for each column. A minimum of 2 valid values in the iPSC and 3 valid values in the iMN samples were required. Imputation was performed for the iPSC samples, but not the iMN samples due to having 5 biological replicates and two technical replicates for the motor neuron data compared to only 3 biological replicates in the iPSC data. Imputation involved replacing missing values from a normal distribution with the default Perseus parameters. Two sample student’s t-tests were performed with a p-value cutoff of 0.05 for significance.

### Antibodies

The following primary antibodies were used for immunoblotting and immunoprecipitation: anti-UBQLN2 (ab190283, Abcam; 85509S, Cell Signaling Technology), anti-BAG6 (sc-365928, Santa Cruz Biotechnology), anti-PEG10 (14412-1-AP, Proteintech), and anti-β-Actin (4967S, Cell Signaling Technology). The following secondary antibodies were used for immunoblotting: IRDye 680RD goat anti-mouse (926-68070) and IRDye 800CW goat anti-rabbit (926-32211), both from LI-COR.

### BAG6 knockdowns

The shRNA lentiviral vector targeting BAG6 was purchased from Sigma (TRCN0000007353). 293FT (Invitrogen, R70007) cells were cultured in Dulbecco’s Modified Eagle Medium (DMEM; Corning, 10-013-CV) supplemented with 10% fetal bovine serum (Atlanta Biologicals) and 1% Penicillin-Streptomycin (Corning, 30-002-CI) and maintained at 37°C in a humidified incubator with 5% CO_2_. Lentiviral particles were produced by transient transfection of 293FT cells with shRNA vector, psPAX2 (Addgene, plasmid #12260) and pCMV-VSV-G (Addgene, plasmid #8454) in a ratio of 6:4:1 as described [41; 42]. Viral supernatants harvested at 24 h and 48 h post-transfection were incubated with iPSCs for 24 h followed by selection in media containing 2 µg/ml puromycin for 72 h.

### Protein extraction, cell fractionation and immunoblotting

For subcellular fractionation, cells were resuspended in a lysis buffer containing 20 mM Tris-HCl (pH 8.0), 138 mM NaCl, 10 mM KCl, 1 mM MgCl_2_, 1 mM EDTA, and 1% CHAPS (vol/vol), supplemented with 20 mM NaF, 20 mM β-glycerophosphate, and protease inhibitor cocktail (Sigma, P8340-5ml). The cells were incubated on ice for 10 min, then centrifuged at 15,000 × g for 10 min at 4°C to separate soluble and insoluble fractions. Western blot images were acquired using the Odyssey XF imaging system (LI-COR Biosciences) and analyzed with ImageStudio software (v5.2, LI-COR Biosciences).

### Proximity-ligation assay (PLA)

iPSCs expressing UBQLN2^WT^, UBQLN2^P497H^, UBQLN2^4XALS^, or UBQLN2^I498X^ were fixed with 4% paraformaldehyde (PFA) and permeabilized with 0.2% Triton X-100 in PBS. The cells were incubated with primary antibodies (α-UBQLN2, 85509S, CST, 1:500; α-BAG6, sc-365928, SC, 1:500) overnight. Reagent kits for Duolink® Proximity Ligation Assay (Sigma) were used, and PLA was performed according to the manufacturers’ conditions. Images were acquired using a Nikon AXR confocal microscope using a 60× oil lens. PLA puncta were quantified using Fiji (ImageJ) automatic particle analysis. Puncta counts were normalized to the number of DAPI-stained nuclei to calculate PLA puncta per cell.

### Statistical Analysis

Statistical analysis information including individual biological and technical replicates number, mean or median, and error bars are explained in the figure legends. Statistical tests were performed in Prism (v10, GraphPad). The tests performed and the resulting *P* values are listed in the figure legends.

## Results

### UBQLN2^ALS^ mutations alter protein-protein interaction networks in iPSCs

To explore how ALS-associated UBQLN2 mutations influence UBQLN2 function through altered protein interactions, we performed label-free quantitative MS on immunoprecipitated UBQLN2 complexes from human iPSCs. We compared wild-type UBQLN2 (UBQLN2^WT^) to three ALS-associated UBQLN2 alleles: a common clinical point mutation (UBQLN2^P497H^), a 4XALS mutant carrying four clinical mutations (UBQLN2^4XALS^: P497H, P506T, P509S, P525S) previously shown to exhibit increased aggregation and toxicity [29; 33], and a truncation mutant (UBQLN2^I498X^) that is undetectable by Western blotting due to its instability [29; 33].

When comparing UBQLN2^WT^ with UBQLN2^I498X^, we identified 24 proteins that were significantly enriched in UBQLN2^WT^ IPs (Fig. 1A, B). As expected, UBQLN2 was the most enriched interactor (~6-fold). Additional interactors included UBQLN1, the ferroptosis-related enzymes VKORC1L1 and VKORC1, and the HSP70 family member HYOU1 (also known as GRP170), a chaperone upregulated by hypoxia. The presence of these interactors is consistent with prior reports of ubiquilin association with chaperone and redox-regulatory proteins [43; 44].

**Figure 1.**
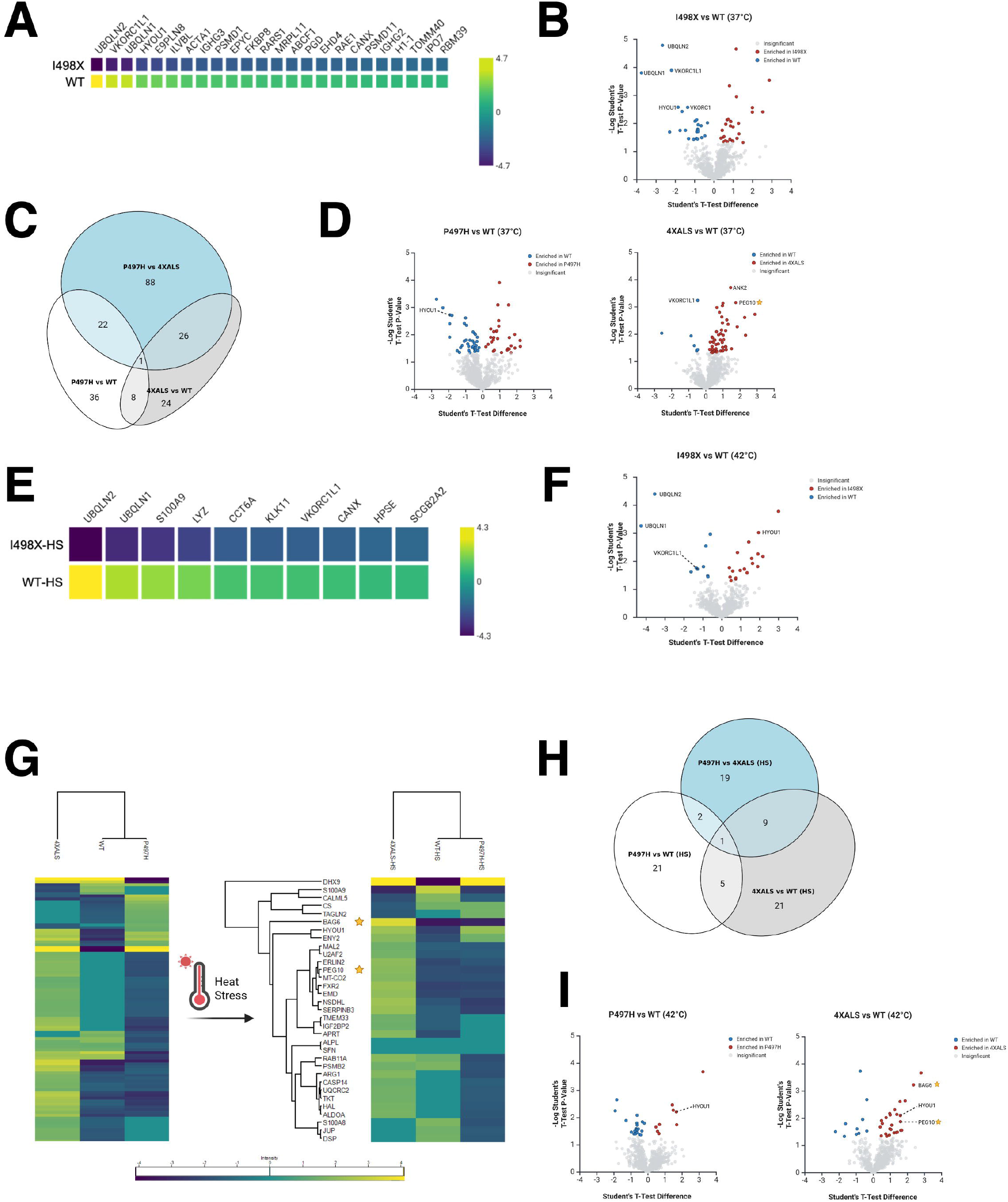
ALS-linked UBQLN2 mutations remodel the UBQLN2 interactome in iPSCs under basal and heat stress conditions. **(A)** Heatmap of proteins differentially enriched in UBQLN2^WT^ and UBQLN2^I498X^ backgrounds under basal conditions. Genes were included if UBQLN2^WT^ mean intensity was greater than UBQLN2^I498X^ mean intensity with p < 0.05. The top 20 genes were selected based on highest WT-UBQLN2 expression. The color scale indicates relative intensity across samples. **(B)** Volcano plot showing proteins differentially enriched in UBQLN2^WT^ versus UBQLN2^I498X^ IPs under basal conditions. Selected interactors (e.g., UBQLN1, VKORC1L1, HYOU1) are highlighted. **(C)** Venn diagram illustrating the overlap and divergence of differentially associated proteins across UBQLN2 variants. Most interactors are uniquely altered in UBQLN2^4XALS^. **(D)** Volcano plot showing proteins differentially enriched in UBQLN2^WT^ versus UBQLN2^P497H^ or UBQLN2^WT^ versus UBQLN2^4XALS^ IPs under basal conditions. Selected interactors (e.g., VKORC1L1, HYOU1, PEG10, ANK2) are highlighted. **(E)** Heatmap of proteins differentially enriched in UBQLN2^WT^ and UBQLN2^I498X^ backgrounds under HS. **(F)** Volcano plot showing proteins differentially enriched in UBQLN2^WT^ versus UBQLN2^I498X^ IPs under HS. **(G)** Heatmap of proteins differentially enriched across UBQLN2^WT^, UBQLN2^P497H^, and UBQLN2^4XALS^ backgrounds under basal and HS conditions. PEG10 and BAG6 are selectively enriched in the 4XALS mutant. **(H)** Venn diagram illustrating the overlap and divergence of differentially associated proteins across UBQLN2 variants under HS. **(I)** Volcano plot showing proteins differentially enriched in UBQLN2^WT^ versus UBQLN2^P497H^ or UBQLN2^WT^ versus UBQLN2^4XALS^ IPs under HS.

We next examined differences in interactome profiles across ALS-associated UBQLN2 alleles. Compared to UBQLN2^WT^, UBQLN2^P497H^ showed differential enrichment of 67 proteins (30 enriched, 37 depleted), while UBQLN2^4XALS^ showed 59 differentially enriched proteins, of which 53 were uniquely enriched in the UBQLN2^4XALS^ background (Fig. 1C, D). Notably, only four proteins overlapped between the UBQLN2^P497H^ and UBQLN2^4XALS^ datasets, suggesting that the 4XALS mutation elicits broader and more distinct interaction changes. Among the most strongly enriched interactors in the 4XALS IPs was PEG10, a retroelement-derived protein previously identified as a UBQLN2 degradation target with important roles in placental biology [24; 45; 46]. Also enriched was ANK2, a cytoskeletal scaffolding protein. Interestingly, the redox regulator and apoptosis suppressor VKORC1L1 [43] was significantly depleted in UBQLN2^4XALS^ IPs compared to all other UBQLN2 variants, indicating a potential LOF interaction specific to the 4XALS mutant. These findings suggest that while UBQLN2^4XALS^ gains aberrant associations (e.g., PEG10, ANK2), it simultaneously loses physiologic interactions such as with VKORC1L1.

To determine whether proteotoxic stress further modulates the UBQLN2 interactome, we repeated the UBQLN2 IP-MS analysis after exposing UBQLN2^WT^, UBQLN2^P497H^, UBQLN2^4XALS^ and UBQLN2^I498X^ iPSCs to heat stress (HS, 42°C, 1 h). In UBQLN2^WT^ IPs, 42 proteins exhibited significant interaction changes post-HS (10 enriched, 32 depleted), reflecting a stress-induced remodeling of the interactome (Fig. 1E–G). Similarly, UBQLN2^4XALS^ displayed heat stress-responsive changes in 36 proteins relative to UBQLN2^WT^ (Fig. 1 G-I). Notably, PEG10 remained associated with UBQLN2^4XALS^ even after HS. Another significant finding was that HYOU1, which lost interaction with UBQLN2^WT^ after HS, gained association with UBQLN2^4XALS^ under HS, indicating altered recruitment of stress-responsive chaperones (Fig. 1 H, I). Additionally, BAG6— a ubiquitin-binding holdase chaperone involved in triage of mislocalized proteins [47; 48; 49]— was enriched in UBQLN2^4XALS^ versus UBQLN2^WT^ IPs. This finding raised the possibility that BAG6 chaperones misfolded UBQLN2.

### Cell-type-specific remodeling of the UBQLN2 interactome in iMNs

To determine whether UBQLN2 interaction changes extend to neurons, we performed analogous UBQLN2 IP-MS analyses using iMNs expressing wild-type and ALS-mutant UBQLN2 alleles. Pairwise comparisons revealed that 21 proteins were differentially associated between UBQLN2^WT^ and UBQLN2^P497H^, while 28 proteins showed differential enrichment between UBQLN2^WT^ and UBQLN2^4XALS^ (Fig. 2A). Notably, the vast majority of these interactions were enhanced in UBQLN2^4XALS^ IPs, suggesting that this mutant variant acquires aberrant protein associations in iMNs. Similar to what was observed in heat-stressed iPSCs, BAG6 was enriched in UBQLN2^4XALS^ IPs prepared from iMNs, suggesting it is a common ALS-linked interactor (Fig. 2B, C). Other proteins enriched in UBQLN2^4XALS^ IPs included BTBD2, which encodes a Ub E3 ligase involved in cytoskeletal dynamics and Top1-mediated chromatin relaxation [50; 51; 52], and phospholipase B D2 (PLBD2), a lysosomal phospholipase that has been implicated in the lysosomal storage disorder, Batten’s Disease [53]. Conversely, several proteins showed reduced association with UBQLN2^4XALS^, including FAM168A/TCRP1, implicated in Polβ stabilization and base excision repair [54] and BAX, a pro-apoptotic factor involved in mitochondrial stress responses [55] that was deenriched in UBQLN2^P497H^ and UBQLN2^4XALS^ IPs relative to wild-type, suggesting potential LOF interactions shared across ALS alleles. PEG10, which was strongly enriched in UBQLN2^4XALS^ IPs in iPSCs, was not detected in iMN IPs, likely reflecting its low expression level in this cell type. Altogether, these results demonstrate that ALS-associated UBQLN2 mutations—particularly the aggregation-prone 4XALS allele—remodel the UBQLN2 interactome in a cell type- and stress-dependent manner. This remodeling involves both aberrant gains of interaction with stress-responsive proteins (e.g., PEG10, BAG6, HYOU1) and loss of interactions (e.g., VKORC1L1, BAX), pointing to a complex interplay of gain- and loss-of-function mechanisms in UBQLN2-linked ALS.

**Figure 2.**
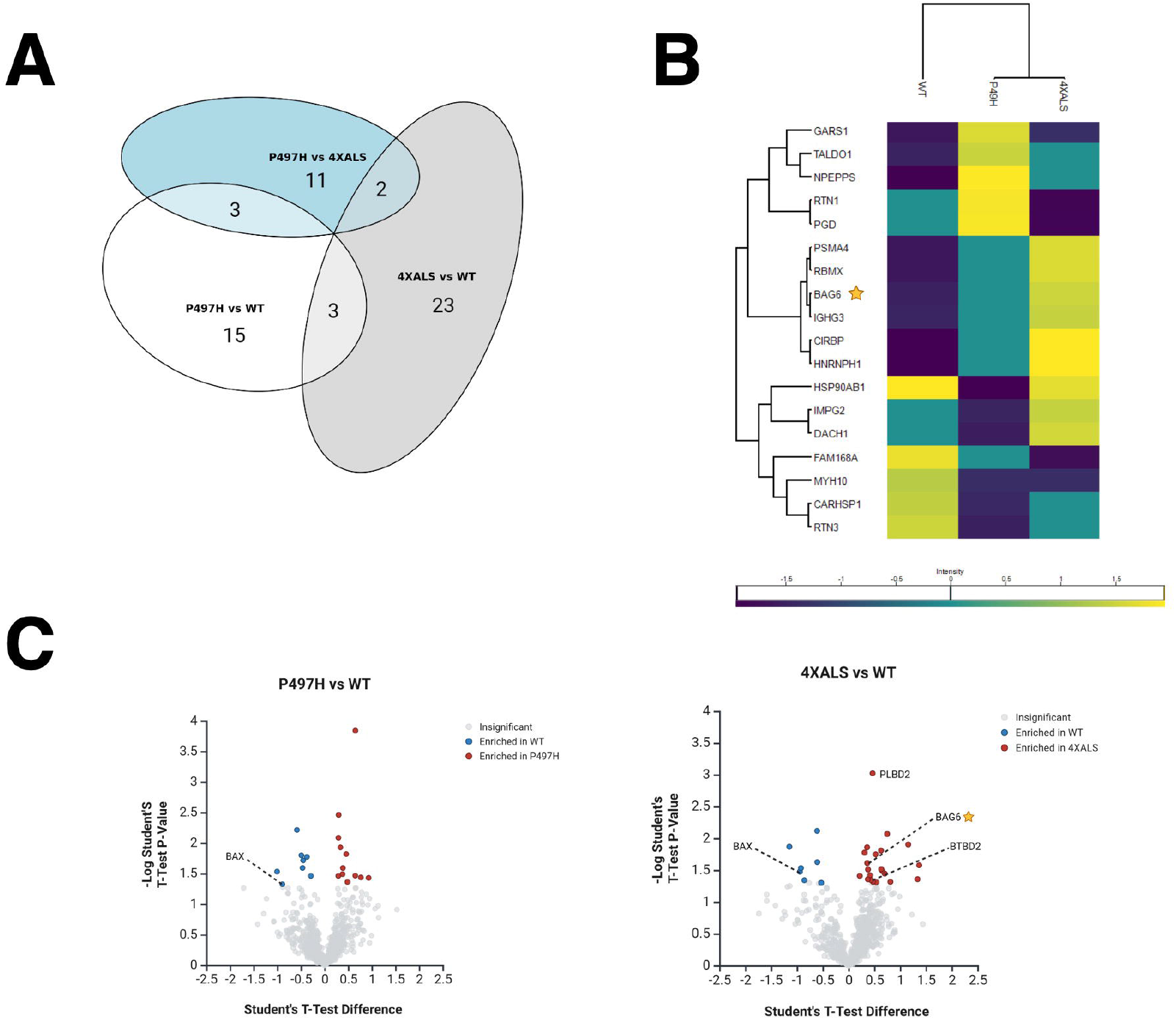
UBQLN2^ALS^ mutations alter protein interaction networks in iMNs. **(A)** Venn diagram illustrating overlap and divergence of differentially associated proteins across the indicated UBQLN2 variants. (**B)** Heatmap showing proteins differentially associated with UBQLN2^WT^, UBQLN2^P497H^, and UBQLN2^4XALS^ in iMNs. BAG6 and PLBD2 are enriched in 4XALS IPs; BAX is depleted. **(C)** Volcano plot showing proteins differentially associated with UBQLN2^WT^, UBQLN2^P497H^, and UBQLN2^4XALS^ in iMNs.

### UBQLN2^4XALS^ exhibits enhanced interaction with BAG6

Given the consistent enrichment of BAG6 in UBQLN2^4XALS^ IP from both iPSCs and iMNs (Fig. 1G, 2C), we investigated this interaction more closely. BAG6 is a multifunctional holdase chaperone best known for its role in the guided entry of tail-anchored (GET) protein pathway, where it facilitates the insertion of tail-anchored (TA) proteins into the endoplasmic reticulum (ER), nuclear, and mitochondrial membranes [47; 56; 57]. In this context, BAG6 forms a cytosolic complex with GET4 (TRC35) and GET5 (UBL4A) to mediate the transfer of TA proteins from the ribosome-associated co-chaperone SGTA to GET3, which then targets them to the GET1–GET2 membrane receptor complex. Outside the canonical GET pathway, BAG6 also plays an important role in triaging MLPs by recognizing exposed hydrophobic regions and directing aberrant proteins for degradation [58].

To validate these MS findings, we performed co-IP followed by Western blotting in UBQLN2 iPSCs. These experiments confirmed that UBQLN2^4XALS^ exhibits a markedly stronger interaction with BAG6 compared to UBQLN2^WT^ (Fig. 3A). To further confirm this association in situ, we employed a proximity ligation assay (PLA), which also revealed significantly increased UBQLN2– BAG6 interaction in UBQLN2^4XALS^ cells relative UBQLN2^WT^ or UBQLN2^P497H^ cells (Fig. 3B, C). These data support BAG6 as a physiologically relevant interactor of ALS-associated UBQLN2 mutants.

**Figure 3.**
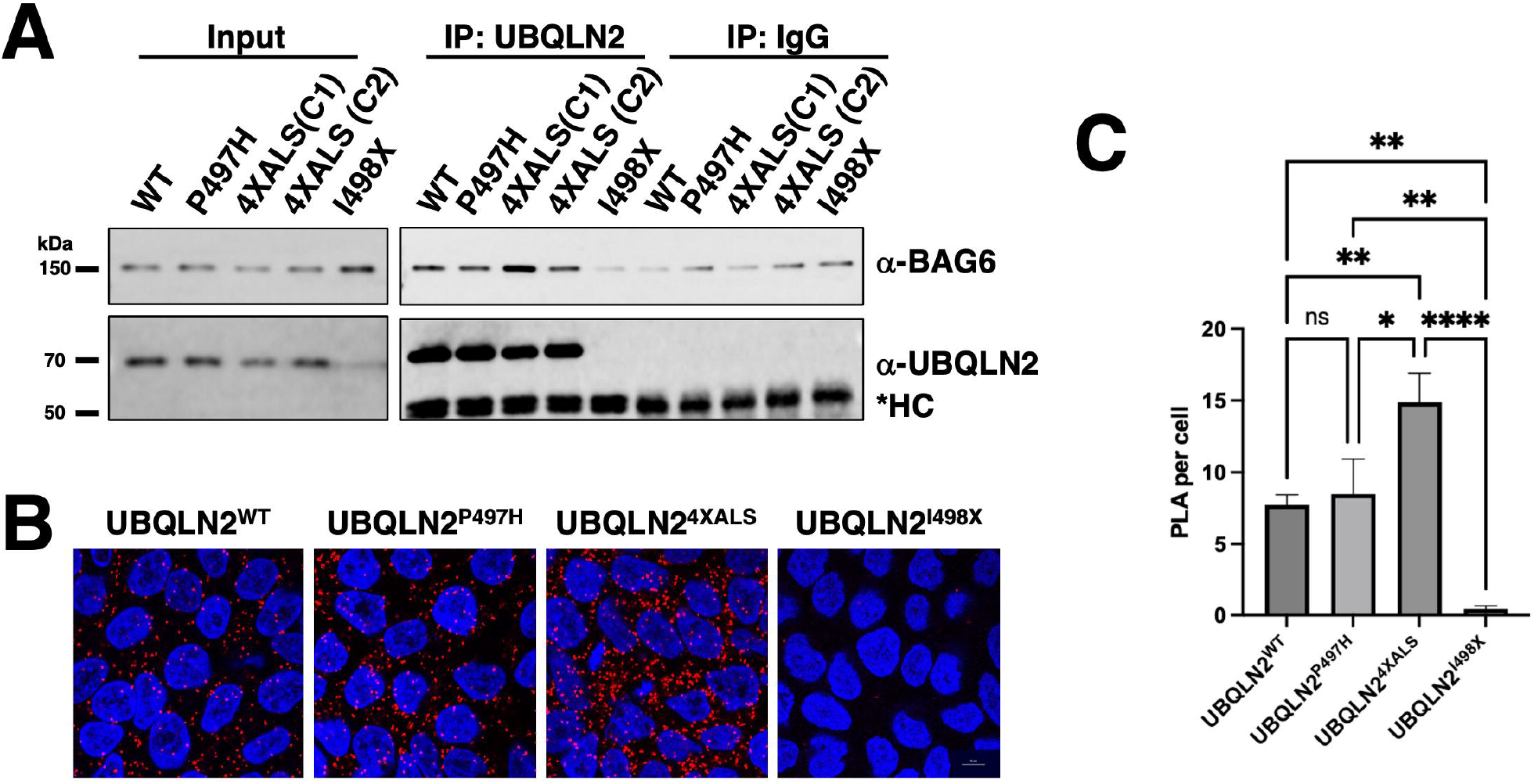
UBQLN2^4XALS^ shows enhanced interaction with BAG6. **(A)** α-UBQLN2 IPs prepared from UBQLN2^WT^, UBQLN2^P497H^, UBQLN2^4XALS^ (two independent clones, C1 and C2), and UBQLN2^I498X^ iPSCs were Western blotted with α-BAG6 antibodies, confirming increased UBQLN2–BAG6 interaction in UBQLN2^4XALS^ cells. **(B)** Representative confocal images of UBQLN2^WT^, UBQLN2^P497H^, UBQLN2^4XALS^, and UBQLN2^I498X^ iPSCs following PLA using α-UBQLN2 and α-BAG6 antibodies (red) and DAPI staining (blue). Magnification: 60×. Scale bar: 10 μm. **(C)** Statistical analysis of PLA data was performed using one-way ANOVA with Tukey’s post hoc test. Data are presented as mean ± SEM from >20 randomly selected regions across three independent experiments; *p ≤ 0.05, **p ≤ 0.01, ****p ≤ 0.001.

### Effect of BAG6 silencing on UBQLN2 solubility

We next investigated whether BAG6 modulates UBQLN2 proteostasis by assessing protein solubility under HS. iPSCs expressing UBQLN2^WT^, UBQLN2^P497H^, or UBQLN2^4XALS^, along with either control or BAG6-targeting shRNAs, were subjected to a 42°C HS for 2 h followed by a 2 h recovery period. Under basal conditions, the solubility of all three UBQLN2 variants was comparable (Fig. 4A, B). Following HS, UBQLN2^WT^ and UBQLN2^P497H^ became less soluble, but their solubility partially recovered after 2 h. In contrast, UBQLN2^4XALS^ exhibited a more pronounced loss of solubility that failed to recover during the post-stress period (Fig. 4A, B). These results indicate that the UBQLN2^4XALS^ mutant is more vulnerable to proteotoxic stress and has a diminished capacity to regain solubility. BAG6 knockdown impaired the solubility recovery of UBQLN2^WT^ and UBQLN2^P497H^ following HS but did further reduce the severe solubility defect of UBQLN2^4XALS^ (Fig. 4A-C). There was also a trend toward increased UBQLN2 expression in shBAG6 cells, however, the differences were not statistically significant (Fig. 4C). Taken together, these findings suggest that BAG6 suppresses UBQLN2 misfolding under conditions of HS, but that its endogenous activity is insufficient to fully rescue the HS-induced aggregation of the UBQLN2^4XALS^ mutant.

**Figure 4.**
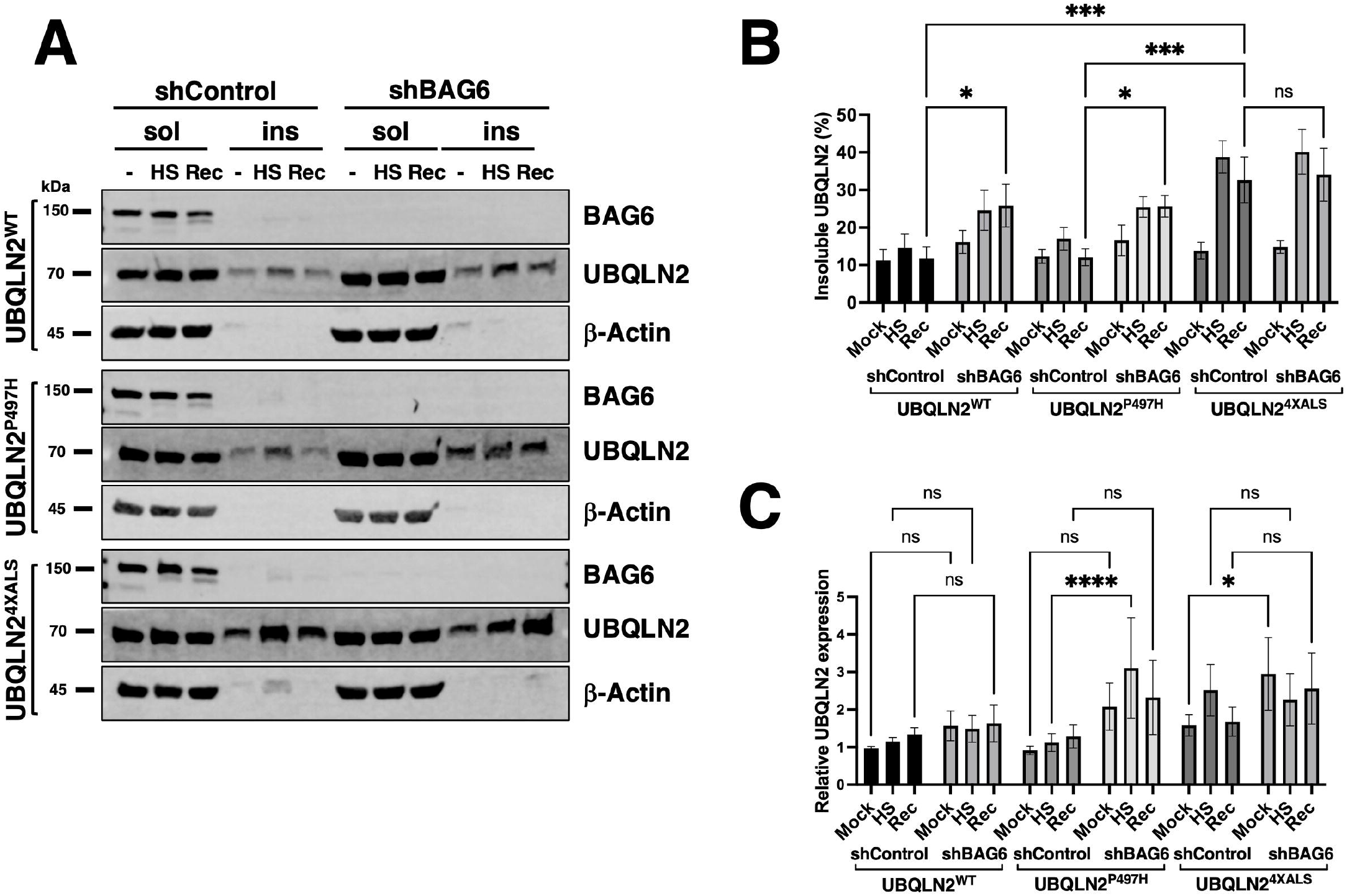
Differential effects of BAG6 silencing on UBQLN2 solubility recovery after HS. **(A)** iPSCs of the indicated UBQLN2 genotypes expressing control or BAG6 shRNA were harvested before, immediately after, or 2 h after exposure to HS (42°C, 2h) and detergent soluble and insoluble fractions analyzed by Western blotting using the indicated antibodies. **(B)** Quantification of insoluble UBQLN2 levels as a proportion of total UBQLN2 protein from immunoblots shown in (A). UBQLN2^4XALS^ showed sustained insolubility following HS compared to UBQLN2^WT^ and UBQLN2^P497H^. **(C)** Total UBQLN2 protein levels were unaffected by BAG6 knockdown. Statistical analysis of (B, C) was performed using two-way ANOVA with Fisher’s LSD post hoc test. Data are presented as mean ± SEM from eight independent experiments; *p ≤ 0.05, ***p ≤ 0.005, ****p ≤ 0.001.

### UBQLN2 and BAG6 coregulate PEG10 degradation

An enhanced interaction between PEG10 and UBQLN2^4XALS^ relative to UBQLN2^WT^ was also observed in iPSC MS experiments (Fig. 1D, G, I). *PEG10* is a domesticated retroviral gene that undergoes programmed −1 ribosomal frameshifting after codon 319 to produce two functionally distinct isoforms (Fig. 5A and Ref [59]). PEG10-reading frame 1 (RF1) encodes a ~50 kDa protein containing Zn finger, Gag, and Pol domains while PEG10-RF1/2 produces a ~100 kDa protein notable for the additional presence of a proline-rich region and aspartyl protease domain [59]. Interestingly, BAG6 was previously found to associate with PEG10, raising the possibility that BAG6, UBQLN2, and PEG10 form a ternary complex [45]. To investigate this hypothesis, we measured PEG10-UBQLN2 interactions in UBQLN2^WT^, UBQLN2^P497H^, and UBQLN2^4XALS^ iPSCs in the absence or presence of BAG6 knockdown. The amount of PEG10 that coimmunoprecipitated with UBQLN2 did not significantly differ between UBQLN2^WT^, UBQLN2^P497H^, and UBQLN2^4XALS^ iPSCs expressing a control shRNA (shControl), though there was a trend for higher co-IP of PEG10 with UBQLN2^P497H^ and UBQLN2^4XALS^ (Fig. 5A, B). BAG6 knockdown caused a significant increase in the level of PEG10 that coimmunoprecipitated with UBQLN2^4XALS^ when compared to UBQLN2^4XALS^:shControl, UBQLN2^WT^:shControl, or UBQLN2^WT^:shBAG6 iPSCs. The co-IP between UBQLN2^P497H^ and PEG10 also trended higher in shBAG6 cells, but was not significantly different from UBQLN2^WT^ iPSCs expressing control or BAG6 shRNA (Fig. 5A, B).

**Figure 5.**
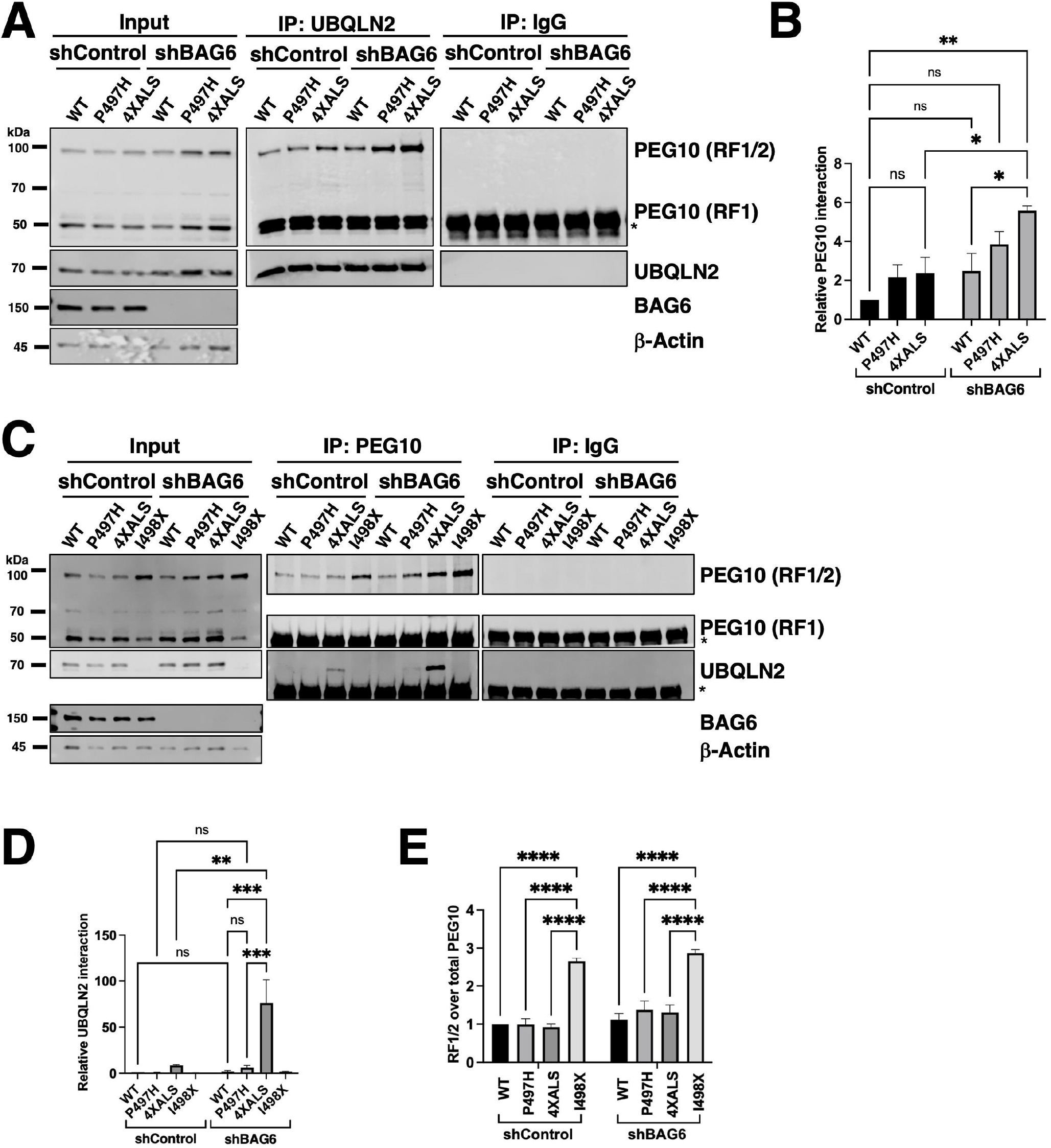
UBQLN2^4XALS^ exhibits enhanced association with PEG10 but retains degradation activity. **(A)** co-IP of UBQLN2 and PEG10 from iPSCs expressing UBQLN2^WT^, UBQLN2^P497H^, or UBQLN2^4XALS^ with or without BAG6 knockdown. Immunoblotting with α-PEG10 antibodies revealed increased UBQLN2–PEG10 interaction in UBQLN2^P497H^ and UBQLN2^4XALS^ backgrounds, particularly under BAG6 knockdown conditions. **(B)** Quantification of PEG10 co-IP across genotypes demonstrates elevated PEG10 binding to UBQLN2^P497H^ and UBQLN2^4XALS^ compared to UBQLN2^WT^. Statistical analysis of (A) was performed using two-way ANOVA with Tukey’s post hoc test. Data are presented as mean ± SEM from three independent experiments; *p ≤ 0.05, **p ≤ 0.01. **(C)** Reciprocal co-IP of UBQLN2 using α-PEG10 antibodies in UBQLN2^WT^, UBQLN2^P497H^, UBQLN2^4XALS^, or UBQLN2^I498X^ iPSCs with or without BAG6 knockdown. **(D)** Quantification of UBQLN2 co-IP results from panel C demonstrated elevated PEG10 binding to UBQLN2^P497H^ and UBQLN2^4XALS^ compared to UBQLN2^WT^ under conditions of BAG6 knockdown. Statistical analysis of (C) was performed using two-way ANOVA with Tukey’s post hoc test. Data are presented as mean ± SEM from three independent experiments; **p ≤ 0.01, ***p ≤ 0.005. **(E)** Quantification of PEG10 RF1/2 levels from input lysates in (C), normalized to total PEG10 (RF1/2+RF1). Data were analyzed by two-way ANOVA with Tukey’s post hoc test. Values represent mean ± SEM from three independent experiments; ****p ≤ 0.0001.

Reciprocal coimmunoprecipitation (co-IP)-WB experiments revealed that levels of immunoprecipitated PEG10 were comparable between UBQLN2^WT^, UBQLN2^P497H^, and UBQLN2^4XALS^ iPSCs but higher in UBQLN2^I498X^ iPSCs, which is consistent with higher expression of PEG10 in UBQLN2-deficient cell lines (Fig. 5C, E) [24; 60]. Similarly, BAG6 knockdown increased PEG10 immunoprecipitation across all UBQLN2 genotypes, with progressively higher levels seen in UBQLN2^WT^, UBQLN2^P497H^, UBQLN2^4XALS^ and UBQLN2^I498X^ iPSCs (Fig. 5C, D). Levels of coimmunoprecipitated UBQLN2 were significantly higher in PEG10 IPs prepared from UBQLN2^4XALS^:shBAG6 iPSCs versus UBQLN2^4XALS^:shControl iPSCs. The co-IP of UBQLN2^WT^ with PEG10 antibodies was not detected either in the absence or presence of BAG6 knockdown while co-IP of UBQLN2^P497H^ was only detectable in UBQLN2^P497H^: shBAG6 iPSCs (Fig. 5C, D). PEG10 expression levels were slightly increased in UBQLN2^P497H^:shBAG6 and UBQLN2^4XALS^:shBAG6 iPSCs relative to UBQLN2^WT^:shBAG6 iPSCs, which may contribute to the enhanced UBQLN2-PEG10 co-IP seen in these lines.

We also noted that the relative abundance of PEG10 isoforms—RF1 (short) and RF1/2 (long)— was differentially affected by UBQLN2 and BAG6 status. UBQLN2^I498X^ iPSCs exhibited a selective increase in the longer RF1/2 isoform, resulting in an increased RF1/2 to RF1 ratio while shBAG6 cells exhibited increases in both RF1 and RF1/2 species and an unchanged RF1/2 to RF1 ratio (Fig. 5E). The isoform shift to RF1/2 persisted in cells with both UBQLN2 deletion and BAG6 knockdown despite the overall increase in total PEG10 levels (Fig 5A, B). These results indicate that pan-PEG10 abundance is primarily controlled by BAG6, while UBQLN2 more specifically regulates PEG10 RF1/2 abundance.

### Reduced PEG10 degradation rate in UBQLN2^4XALS^ iPSCs

UBQLN2^P497H^ and UBQLN2^4XALS^ alleles showed increased interaction with PEG10 in the setting of BAG6 deficiency, and this effect was most pronounced for UBQLN2^4XALS^ (Fig. 5C). While steady state levels of PEG10 were not as upregulated in UBQLN2^4XALS^:shBAG6 iPSCs as they were in UBQLN2^I498X^:shBAG6 iPSCs, this finding raised the possibility that UBQLN2^4XALS^ is defective for PEG10 degradation. To more closely examine this possibility, we measured the degradation rate of PEG10 in UBQLN2^WT^, UBQLN2^P497H^, UBQLN2^4XALS^ and UBQLN2^I498X^ iPSCs treated with the protein synthesis inhibitor, cycloheximide (CHX). The levels of PEG10-RF1/2 and PEG10-RF1 decreased with similar kinetics in CHX-treated UBQLN2^WT^ (Fig. 6A) and UBQLN2^P497H^ (Fig. 6B) iPSCs but failed to significantly decrease in UBQLN2^I498X^ (Fig. 6D). UBQLN2^4XALS^ (Fig. 6C) iPSCs exhibited a PEG10 degradation rate intermediate between UBQLN2^WT^ and UBQLN2^I498X^ while BAG6 knockdown increased steady state levels of PEG10 across all UBQLN2 genotypes but did/did not affect PEG10 degradation rate (Fig. 6). These finding suggest that UBQLN2^4XALS^ is a partial LOF allele with respect to PEG10 degradation.

**Figure 6.**
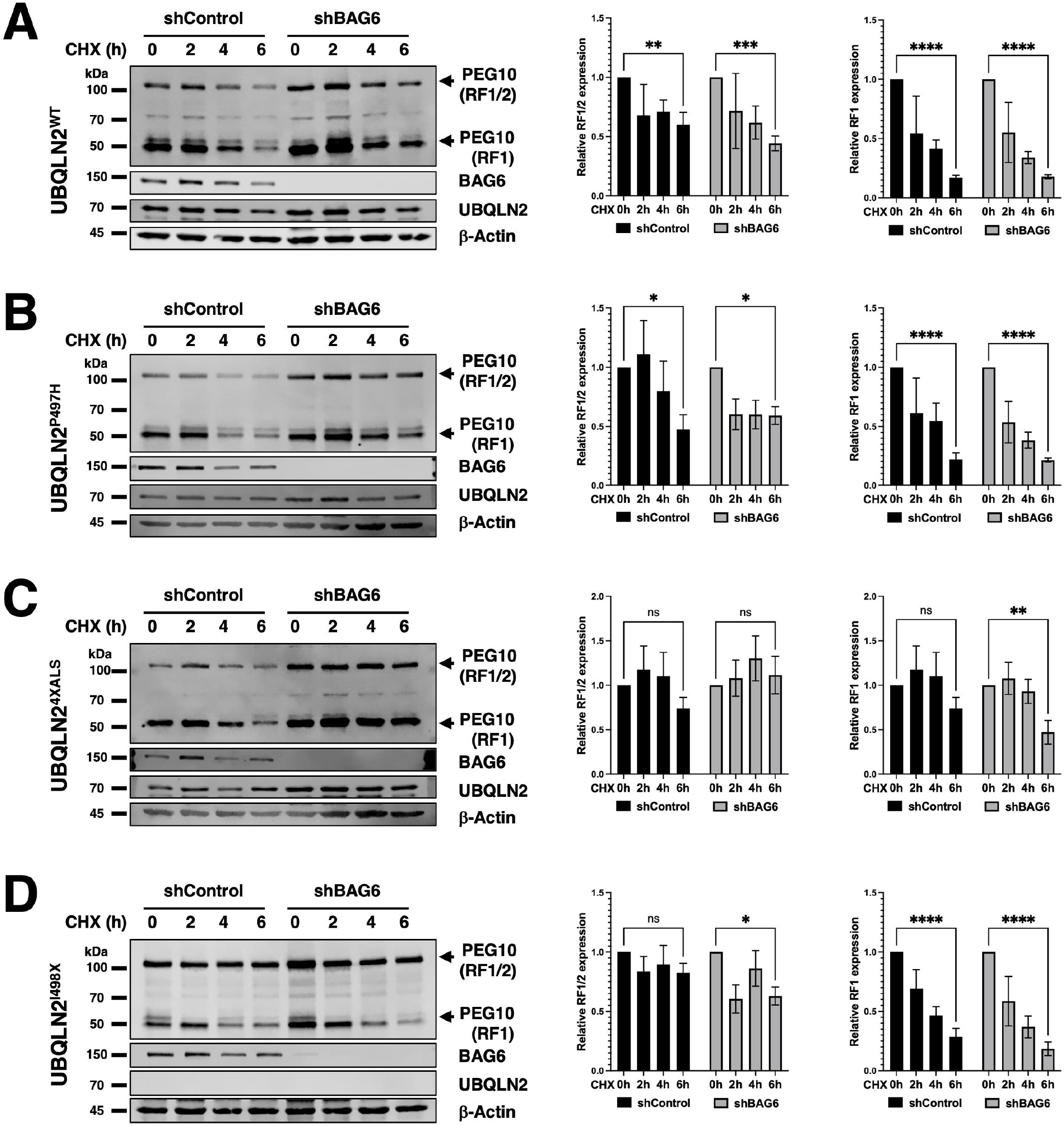
Distinct roles for UBQLN2 and BAG6 in regulating PEG10 isoform degradation. **(A-D)** Western blot analysis of PEG10 isoforms in UBQLN2^WT^ (A), UBQLN2^P497H^ (B), UBQLN2^4XALS^ (C), or UBQLN2^I498X^ (D) iPSCs with or without BAG6 knockdown following treatment with 50 μg/ml CHX for the indicated times. Quantification of PEG10 RF1 and RF1/2 isoform levels in Western blots normalized to loading control β-actin, shows degradation rate of RF1/2 and RF1 in iPSCs. Data were analyzed by two-way ANOVA with Tukey’s post hoc test. Values represent mean ± SEM from three independent experiments; *p ≤ 0.05, **p ≤ 0.01, ****p ≤ 0.0001.

### HS-dependent degradation of PEG10 occurs independent of UBQLN2 and BAG6

Given that BAG6 silencing increased PEG10 levels and diminished UBQLN2 solubility following HS, we examined whether UBQLN2 mutations or BAG6 expression influenced PEG10 expression and solubility under HS conditions. Levels of PEG10-RF1 and PEG10-RF1/2 declined in UBQLN2^WT^, UBQLN2^P497H^, and UBQLN2^4XALS^ iPSCs following a 42°C, 2 h HS, with little change in PEG10 solubility (Fig. 7A, B). PEG10 expression recovered to comparable levels in UBQLN2^WT^ and UBQLN2^4XALS^ iPSC within 2 h after retransfer to 37°C, suggesting active resynthesis and/or stabilization (Fig. 7C, D). Similarly, while BAG6 knockdown increased steady-state PEG10 levels, particularly in UBQLN2^4XALS^ iPSCs, it did not affect HS-dependent PEG10 degradation or post-HS recovery (Fig. 7C, D). HS-dependent degradation of PEG10 also occurred in UBQLN2^I498X^ iPSCs, suggesting that neither ALS-associated mutations nor UBQLN2 functional deficiency significantly affect HS-dependent PEG10 degradation (Fig. 7E, F). Thus, HS-dependent PEG10 turnover occurs through a UBQLN2 and BAG6-independent mechanism.

**Figure 7.**
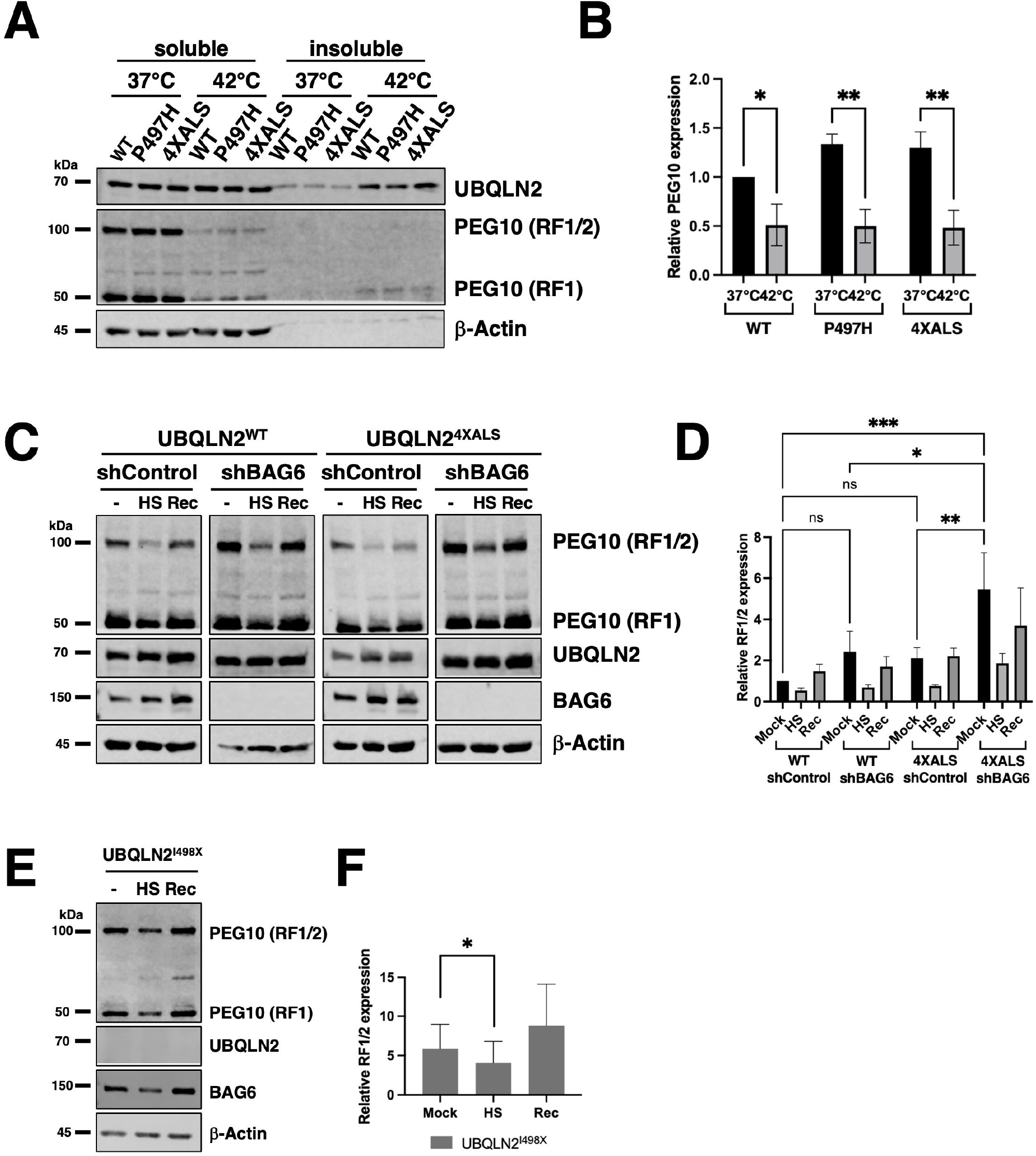
HS induces PEG10 degradation in a UBQLN2- and BAG6-independent manner. **(A)** Western blot analysis of PEG10 RF1 and RF1/2 isoforms in UBQLN2^WT^, UBQLN2^P497H^, or UBQLN2^4XALS^ iPSCs in the absence of HS (37°C) or immediately after HS at 42°C, for 2 h. Cell extracts were separated into soluble and insoluble fractions and analyzed by Western blotting with the indicated antibodies. **(B)** Quantification of PEG10 RF1 and RF1/2 isoforms shows HS– induced degradation across UBQLN2 genotypes. Data were analyzed by two-way ANOVA with Tukey’s post hoc test. Values represent mean ± SEM from three independent experiments; *p ≤ 0.05, **p ≤ 0.01. **(C)** Western blot analysis of PEG10 RF1 and RF1/2 in UBQLN2^WT^ and UBQLN2^4XALS^ iPSCs in the absence or presence of BAG6 knockdown. Cells were harvested in the absence of HS (-); immediately after HS (HS); or 2 h following recovery from HS (Rec) and subjected to fraction and Western blotting with the indicated antibodies. **(D)** Quantification of PEG10 RF1/2 Western blotting data from panel C. Data were analyzed by two-way ANOVA with Tukey’s post hoc test. Values represent mean ± SEM from three independent experiments; *p ≤ 0.05, **p ≤ 0.01. **(E)** HS-dependent degradation of PEG10 in UBQLN2^I498X^ iPSCs, **(F)** Quantification of PEG10 RF1/2 Western blotting data from panel E. Data were analyzed by one-way ANOVA with Tukey’s post hoc test. Values represent mean ± SEM from three independent experiments; *p ≤ 0.05.

## Discussion

UBQLN2 functions at the intersection of protein quality control and degradation, shuttling ubiquitinated substrates to the proteasome and engaging misfolded proteins via stress-responsive chaperone networks [61]. Here, we demonstrated that ALS-associated mutations in the UBQLN2 PRR exert GOF and LOF effects on protein-protein interactions in both iPSCs and iMNs. The gain and loss of relevant protein-protein interactions may contribute to cellular defects that trigger neurodegeneration in ALS/FTD-UBQLN2.

We previously reported that the combinatorial UBQLN2^4XALS^ allele exhibited heightened misfolding and toxicity profiles versus UBQLN2^ALS^ clinical alleles, making this allele a valuable tool for probing UBQLN2 pathomechanisms [29; 33]. Consistent with its more severe folding defect, UBQLN2^4XALS^ mutant showed pronounced changes in its interactome, with a propensity toward gained interactions that were highlighted by increased binding to BAG6, a holdase, and PEG10, a previously described degradation target of UBQLN2 [24; 60].

The enhanced interaction between BAG6 and UBQLN2^4XALS^ is most consistent with a role for BAG6 in the triage of misfolded UBQLN2. BAG6 plays an important role in triaging MLPs by binding to exposed hydrophobic regions, thereby reducing the rate of aberrant intra- and intermolecular interactions that would lead to protein aggregation [58]. We speculate that ALS-associated mutations in the PRR cause exposure of hydrophobic regions, including the Met-rich, central region, leading to recognition by BAG6, with the more severely misfolded UBQLN2^4XALS^ allele, eliciting greater BAG6 engagement. Functional importance of the BAG6-UBQLN2 interaction was suggested by the finding that BAG6 knockdown decreased the solubility of both UBQLN2^WT^ and UBQLN2^P497H^ proteins following recovery of iPSCs from HS (Fig. 4A, B). Ironically, despite increased binding, BAG6 knockdown did not exacerbate the post-HS insolubility phenotype of UBQLN2^4XALS^, suggesting that its insolubility phenotype is saturated (Fig. 4A, B). We also noted that UBQLN2 protein levels were slightly upregulated in shBAG6 iPSCs, suggesting that a fraction of BAG6-bound UBQLN2 is destined for degradation, as seen for other BAG6 clients [56]. Because BAG6 also suppressed aggregation of C-terminal fragments of TDP-43 [62], it may be of broad relevance to understanding pathologic protein misfolding in ALS/FTD.

A second finding of this study is that UBQLN2^ALS^ mutants exhibited increased interaction with PEG10, a degradation target of UBQLN2 [24; 45; 46; 60]. Although, the biochemical basis for the enhanced association between UBQLN2^ALS^ mutants and PEG10 is not clear, it may involve stabilized interactions between ubiquitylated PEG10 and the UBQLN2 UBA domain and/or aberrant association of PEG10 with exposed hydrophobic patches that are also engaged by BAG6. The fact that knockdown of BAG6 potentiated the co-IP of PEG10 with UBQLN2^P497H^ and UBQLN2^4XALS^, but not UBQLN2^WT^ (Fig. 5A-C), is consistent with the idea that misfolded UBQLN2 species form stabilized complexes with PEG10. However, because BAG6 silencing marginally increased the levels of UBQLN2 and significantly increased PEG10 expression (Fig. 4C, 7C, 7D), the increased co-IP stoichiometry between PEG10 and UBQLN2^ALS^ mutants in BAG6 knockdown cells may be partially attributed to increased input of the target proteins. The enhanced co-IP between UBQLN2 and PEG10 in shBAG6 cells also ruled out that BAG6 bridged the PEG10-UBQLN2 complex.

A previous study identified BAG6 as a PEG10-associated protein [45]. Our findings extend this observation to show that BAG6 inhibits PEG10 expression (Fig. 5). However, while both UBQLN2 and BAG6 suppressed PEG10 protein levels; they appear to do so independently and via different mechanisms. Interestingly, UBQLN2 LOF selectively stabilized the RF1/2 PEG10 translation product (Fig. 5C, E, 6D) [24], while BAG6 knockdown stabilized both RF1 and RF1/2 PEG10 species (Fig. 5C, 6). One speculative possibility is that BAG6 regulates PEG10 stability through recognition elements in the PEG10 RF1 while UBQLN2 controls RF1/2 stability by binding to elements unique to RF1/2, including the Pro-rich region and peptidase domains. It is also possible that UBQLN2 inhibits the −1 frame shift required to generate PEG10 RF1/2. Finally, the existence of a third PEG10 degradation pathway is suggested by the fact that neither UBQLN2 nor BAG6 LOF affected HS-dependent PEG10 degradation (Fig. 7C, D). Further studies are needed to discern unique and shared functions of UBQLN2 and BAG6 in controlling PEG10 isoform abundance.

Prior studies reported variable effects of UBQLN2^ALS^ mutants on PEG10 stability [24]. Steady-state expression and half-life of PEG10 were marginally increased in UBQLN2^4XALS^ iPSCs and unchanged in UBQLN2^P497H^ iPSCs relative to UBQLN2^WT^ controls (Fig. 6A-C). By contrast, the UBQLN2^I498X^ mutation had a strong stabilizing effect on PEG10 (Fig. 6D), which is consistent with previous studies [24; 60]. From this, we surmise that the ALS mutations have a modest, quantitative impact on PEG10 stability that correlated with the strength of UBQLN2-PEG10 interaction in co-IP assays. Our findings further indicate that the effects of ALS mutations—and the 4XALS mutation in particular—on PEG10 proteostasis are exacerbated by BAG6 functional insufficiency. This finding may hold relevance to understanding how age-dependent declines in proteostasis synergize with disease mutations to instigate neurodegeneration in ALS/FTD and other NDs [1].

Although our validation studies were primarily focused on BAG6 and PEG10, other proteins showing differential association with UBQLN2^ALS^ mutants may be relevant to understanding the UBQLN2 pathomechanism. In iPSCs, the ER resident HSP70 family member HYOU1 showed reduced association with UBQLN2^ALS^ mutants in iPSCs the absence of HS and increased association with UBQLN2^ALS^ mutants following HS (Fig. 1D, I). Several prior studies have reported that ALS-associated mutations in UBQLN2 reduce its association with canonical HSP70 [44; 63; 64; 65]. Enhanced association between UBQLN2^4XALS^ and the Ub E3 ligase BTBD2 in iMNs may reflect a role for BTBD2 in UBQLN2 turnover while reduced association of UBQLN2^4XALS^ with FAM168A may impact FAM168A-dependent DNA Polβ stabilization and base-excision repair (BER) critical to neuronal integrity [54]. Further studies are needed to define which altered protein-protein interactions are most central to the UBQLN2 pathomechanism in neurons.

## Data Availability Statement

All raw mass spectrometry files and search results were deposited in the MassIVE Repository (MassIVE ID: MSV000099268).

## Author Contributions

RST and SHK contributed to the conception and design of the study. SHK, MS, CEB, and AKW contributed to the acquisition of data. SHK and CEB performed the statistical analysis. RST, SHK, MS, and CEB interpreted the data. RST and SHK wrote the first draft of the manuscript. MS and CEB wrote sections of the manuscript. All authors contributed to manuscript revision, read, and approved the submitted version.

## Generative AI statement

The authors declare that no Generative AI was used in the creation of this manuscript.

## Abbreviations

ALS: amyotrophic lateral sclerosis
FTD: frontotemporal dementia
UBQLN2: Ubiquilin-2
BAG6: BCL2-associated athanogene 6
PEG10: paternally expressed gene 10
iMN: induced motor neuron
HS: heat stress
CHX: cycloheximide.

